## Supplementary Figures 1-10, Supplementary Tables 1-2 for "Unscrambling Fluorophore Blinking for Comprehensive Cluster Detection via Photoactivated Localization Microscopy"

#### Supplementary Figure 1

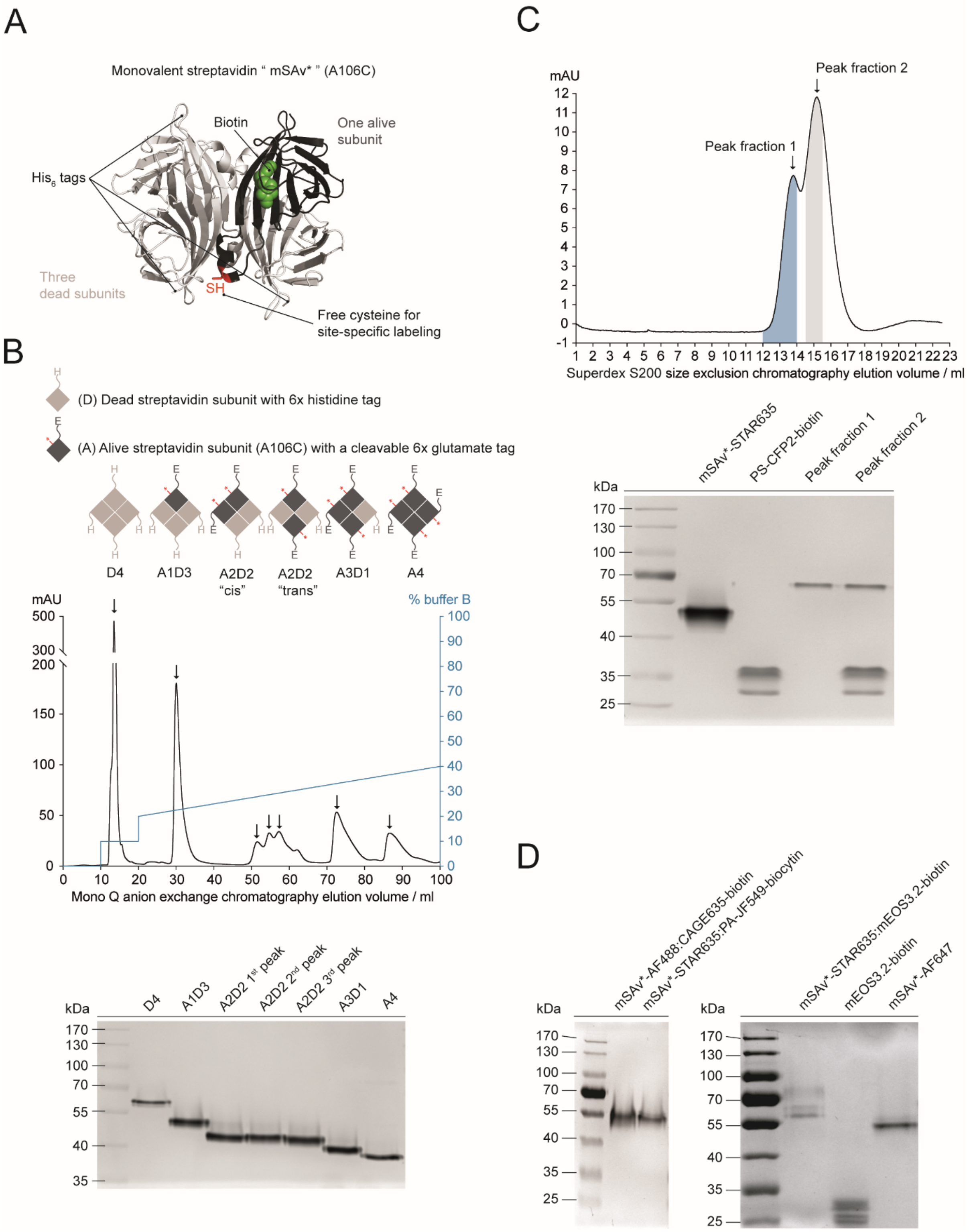


**Supplementary Figure 1: Synthesis of monovalent 3x $His_6$ tagged streptavidin, site-specifically labeled with Abberior STAR635 (or Alexa Fluor 488) and conjugated to biotinylated PS-CFP2, mEOS3.2, PA-JF549 or CAGE635.**

(**A**) Positions for molecular modifications to anchor photoswitchable fluorescent proteins (PS-CFP2 and mEOS3.2) or photoactivatable organic dyes (PA-JF549 and CAGE635) were chosen based on the structure of monovalent streptavidin (adapted from PDB 1SWE).

(**B**) Monovalent streptavidin (A1D3) could be separated from zero-valent (D4), divalent (A2D2), trivalent (A3D1) or tetravalent (A4) streptavidin by means of the C-terminal hexa-glutamate tag located on the "alive" subunit. Major peak fractions from Mono Q anion exchange chromatography were analyzed by SDS PAGE (silver stain) conducted under reducing and non-boiling conditions. Streptavidin tetramers migrate more rapidly due to an increase in negative charge contributed by the hexa-glutamate tags. (A4 > D4).

(**C**) S200 size exclusion chromatography was applied to separate the mSAv*-STAR635:PS-CFP2-biotin conjugate from free PS-CFP2-biotin. SDS-PAGE performed under non-reducing and non-boiling conditions followed by silver staining of the protein bands reveal the protein purity of peak fraction 1.

(**D**) S200 size exclusion chromatography was applied to separate the mSAv*-STAR635:PA-JF549-biocytin, the mSAv*-STAR488:CAGE635-biotin or the mSAv*-STAR635:mEOS3.2-biotin conjugate from free PA-JF549-biocytin, CAGE635-biotin or mEOS3.2-biotin, respectively. SDS-PAGE performed under non-reducing and non-boiling conditions and visualization of the protein bands by silver staining (left) or colloidal Coomassie Brilliant Blue G-250 staining (right) reveal the purity of the indicated protein conjugates.

### Supplementary Figure 2

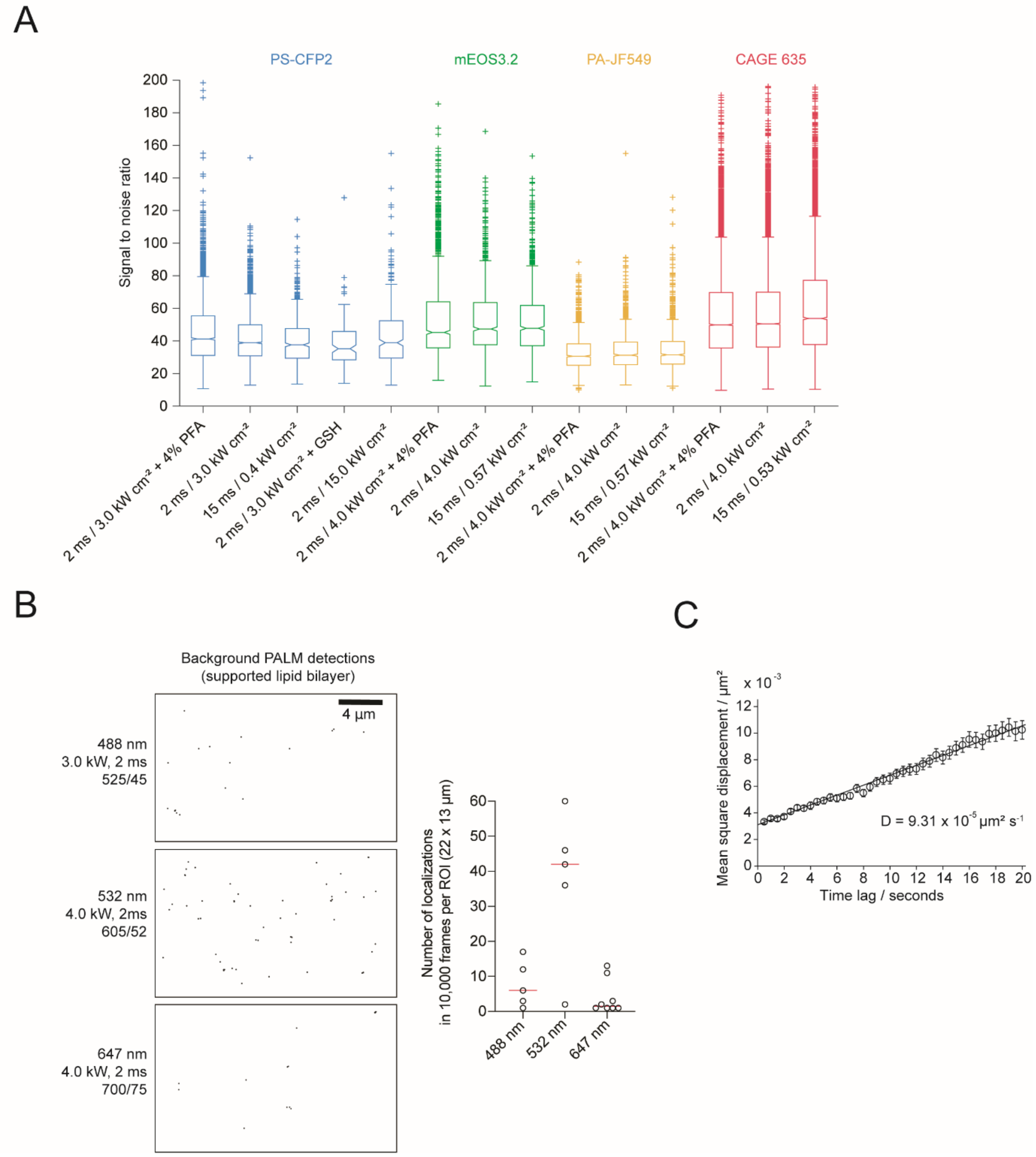


**Supplementary Figure 2: Calculated signal to noise ratios of PS/PA-fluorophores, background signals observed on SLBs during PALM measurements, and single molecule tracking of the SAv platform on DPPC bilayers.**

**(A)** Signal to noise ratios of individual PS-CFP2, mEOS3.2, PA-JF549 and CAGE635 fluorescence signals recorded under the indicated imaging conditions. Ratios were calculated by dividing the integrated brightness of individual signals by the standard deviation of the local background. On each box, the central mark indicates the median, and the bottom and top edges of the box indicate the 25th and 75th percentiles, respectively. Outliers are plotted individually with a '+' symbol (n = 2902 / 3183 / 626 / 142 / 338 PS-CFP2 signals, n = 1280 / 1044 / 1066 mEOS3.2 signals, n = 13387 / 13892 / 7961 CAGE635 signals and n = 1563 / 1113 / 1080 PA-JF549 signals were analyzed).

**(B)** Localization map derived from PALM measurements of 4% PFA-fixed bilayers in the absence of any fluorophores (left) recorded for all color channels employed. The number of localizations detected in 10,000 frames for individual color channels is shown on the right.

(**C**) Single molecule tracking results for mSAv*-STAR635 linked to the DPPC bilayer. The mean square displacement increased linearly with time, characterized by a diffusion constant of $D = 9.31 \times 10^{-5}$ $\mu m^2 s^{-1}$ (n = 79 molecules). Error bars represent SEM.

Supplementary Figure 3

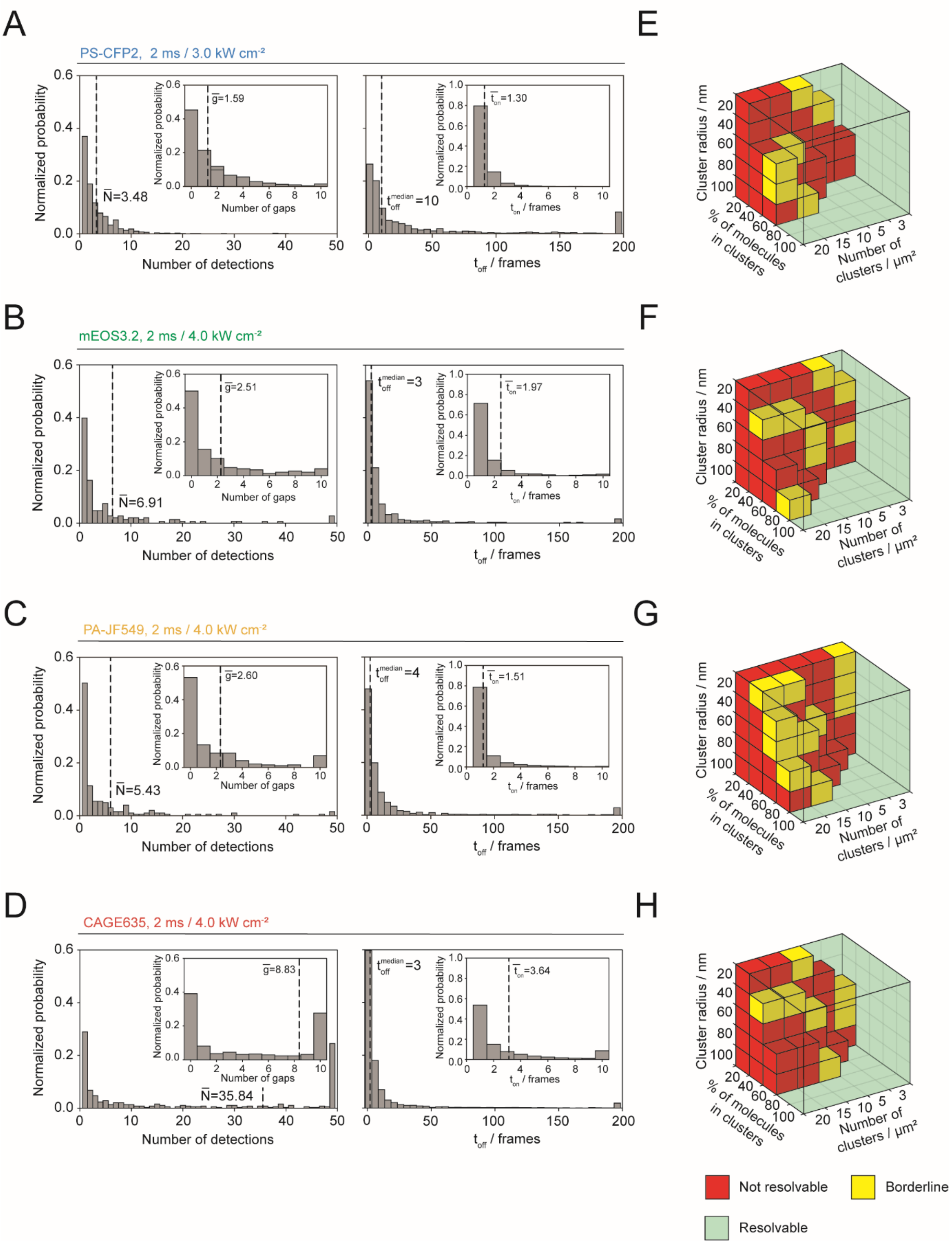


**Supplementary Figure 3: PS-CFP2, mEOS3.2, PA-JF549 and CAGE635 blinking behavior at high power densities and 2 ms illumination.**

(**A-D**) Normalized histograms illustrate four parameters described in (**Fig. 1D**) determined for PS-CFP2, mEOS3.2, PA-JF549 and CAGE635 at the indicated power densities with 2 ms illumination without 4% PFA pretreatment. Numbers within the histograms denote mean or median values of the indicated parameter for 1080 PS-CFP2, 148 mEOS3.2, 203 PA-JF549 and 384 CAGE635 molecules analyzed.

(**E-H**) Sensitivity of Ripley's K analysis in detecting clustering scenarios. Ripley's K functions have been determined through Monte Carlo simulations of nanoclusters with a cluster radius of 20, 40, 60 or 100 nm, a fraction between 20% and 100% of molecules residing inside clusters, 3, 5, 10, 15 and 20 clusters per $\mu m^2$, an average molecular density of 70 molecules $\mu m^{-2}$ and based on blinking statistics of PS-CFP2 (**E**), mEOS3.2 (**F**), PA-JF549 (**G**), and CAGE635 (**H**) determined experimentally under indicated imaging conditions. Functions were compared to those calculated from random distributions and categorized as described in **Fig. 3B**.

#### Supplementary Figure 4

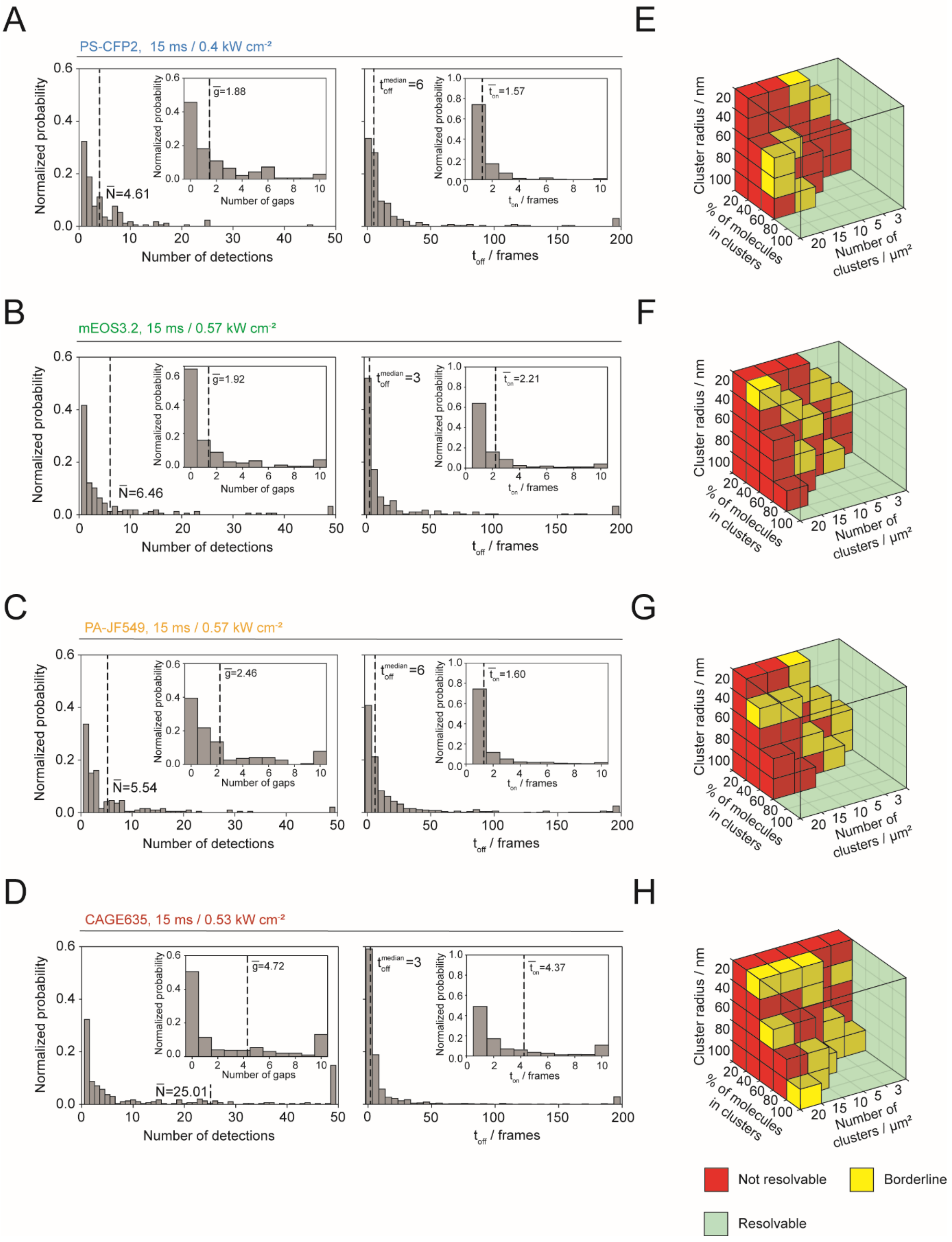


**Supplementary Figure 4: PS-CFP2, mEOS3.2, PA-JF549 and CAGE635 blinking behavior at low power density and 15 ms illumination time.**

(**A-D**) Normalized histograms illustrate four parameters described in (**Fig. 1D**) determined for PS-CFP2, mEOS3.2, PA-JF549 and CAGE635 recorded at the indicated power densities with 15 ms illumination without 4% PFA pretreatment. Numbers within the histograms denote mean or median values of the indicated parameter for 170 PS-CFP2, 166 mEOS3.2, 197 PA-JF549 and 310 CAGE635 molecules analyzed.

(**E-H**) Sensitivity of Ripley's K analysis in detecting clustering scenarios. Ripley's K functions have been determined through Monte Carlo simulations of nanoclusters with a cluster radius of 20, 40, 60 or 100 nm, a fraction between 20% and 100% of molecules residing inside clusters, 3, 5, 10, 15 and 20 clusters per $\mu m^2$, an average molecular density of 70 molecules $\mu m^{-2}$ and based on blinking statistics of PS-CFP2 (**E**), mEOS3.2 (**F**), PA-JF549 (**G**), and CAGE635 (**H**) determined experimentally under indicated imaging conditions. Functions were compared to those calculated from random distributions and categorized as described in **Fig. 3B**.

### Supplementary Figure 5

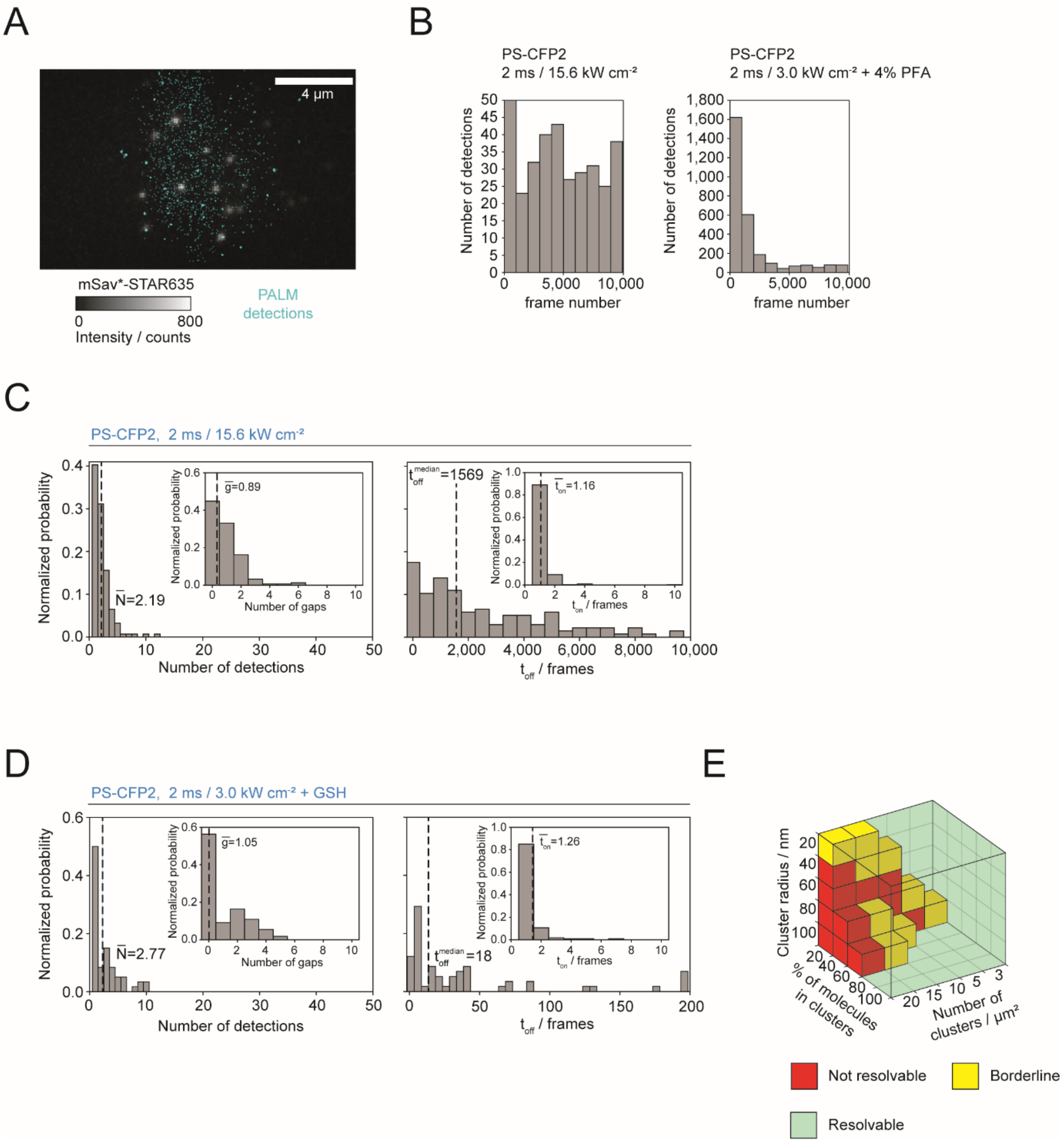


**Supplementary Figure 5: Detailed quantitation of PS-CFP2 blinking behavior at 15 kW/cm² power density or in the presence of reduced glutathione (GSH).**

(**A**) PALM recordings at 15.6 kW $cm^{-2}$ illumination power density were associated with an increase in fluorophore-unrelated localizations. Shown is a two-color overlay of mSAv*-STAR635 (monochrome color channel) recorded prior to PALM with detections from 10,000 consecutive PALM recordings of single PS-CFP2 signals (cyan colored localizations). Nonspecific signals yielded a rather homogeneous background of localizations, interfering with unambiguous identification of true PS-CFP2 signals.

**(B)** Histograms display the number of fluorescence (PS-CFP2) detections per frame recorded at a power density of 15.6 kW $cm^{-2}$ (left) or 3.0 kW $cm^{-2}$ (right) with 2 ms illumination.

**(C, D)** Normalized histograms of four indicated parameters determined for PS-CFP2 recorded at a power density of 15.6 kW $cm^{-2}$ (**C**) and 3.0 kW $cm^{-2}$ with 2 ms illumination in the presence of 5 mM reduced glutathione (GSH) (**D**). Numbers within the histograms denote mean or median values of the indicated parameter determined for 60 recorded molecules.

(**E**) Sensitivity of Ripley's K analysis in detecting clustering scenarios in the presence of reduced glutathione (GSH). Ripley's K functions have been determined through Monte Carlo simulations of nanoclusters with a cluster radius of 20, 40, 60 or 100 nm, a fraction between 20% and 100% of molecules residing inside clusters, 3, 5, 10, 15 and 20 clusters per $\mu m^2$, an average molecular density of 70 PS-CFP2 molecules $\mu m^{-2}$ and based on blinking statistics determined experimentally under indicated imaging conditions. Functions were compared to those calculated from random distributions and categorized as described in **Fig. 3B**.

### Supplementary Figure 6

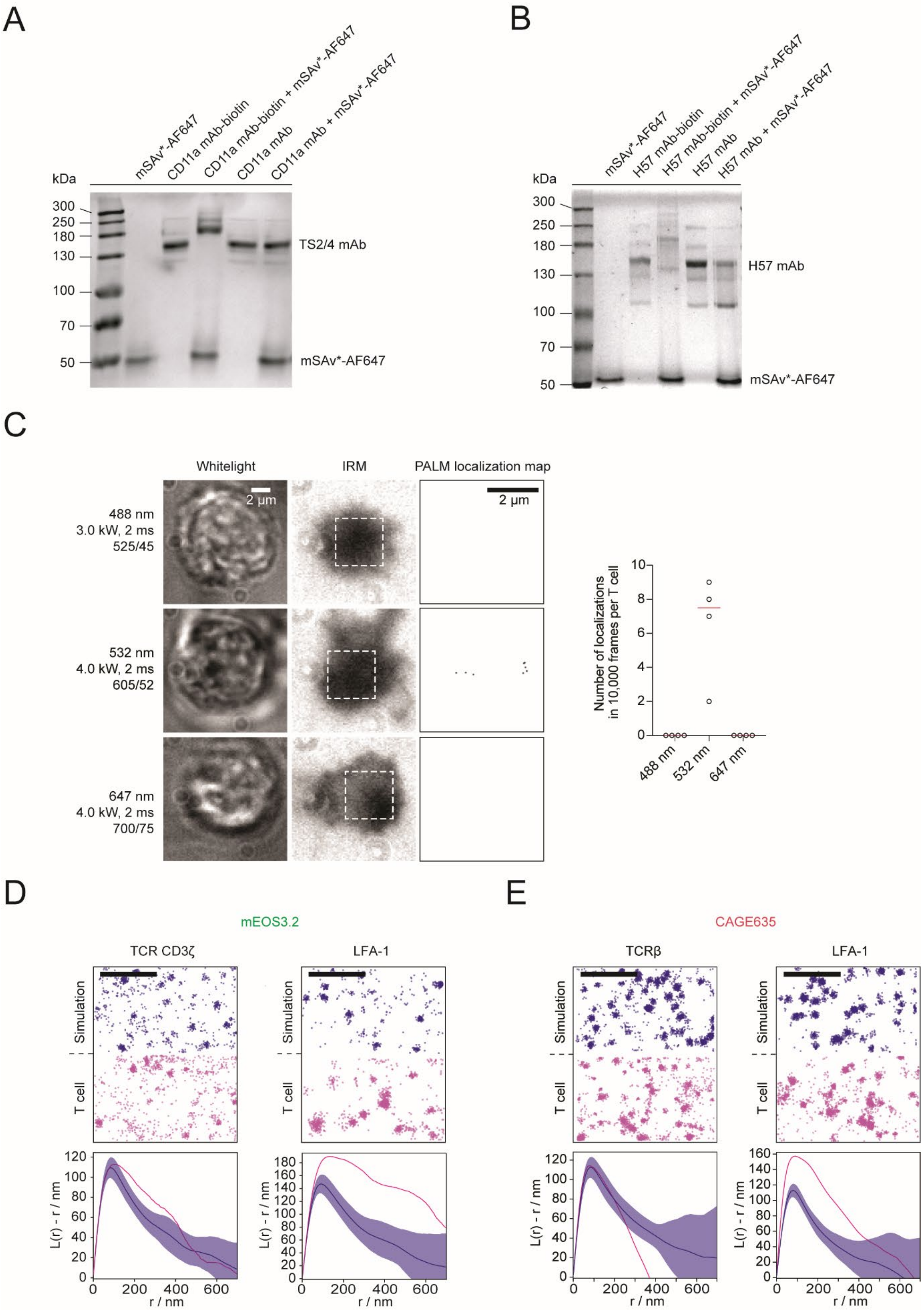

**Supplementary Figure 6: mSAv-based gel shift assays to assess the degree of biotinylated lysine residues on monoclonal antibodies, PALM measurements of cellular background, and classification of CD3ζ- and LFA-1- distributions on PFA-fixed T cells as determined with the use of mEOS3.2 and CAGE635.**

(**A, B**) mSAv-based gel shift assays reveal the degree of biotinylation of the monoclonal antibodies TS2/4 (reactive against LFA-1, CD11a subunit, (**A**)) and H57 (reactive against TCRβ, (**B**)). After purification via immobilized avidin agarose (to remove non-biotinylated mAbs and free NHS-LC-LC-Biotin), and S200 size exclusion chromatography, the biotinylated mAbs were subjected to 8% SDS PAGE analysis in the presence or absence of mSAv. To quantitate the degree of mAb biotinylation, we stained the protein bands with colloidal Coomassie Brilliant Blue G-250. A shift of ~50 kDa, ~100 kDa and 150 kD indicates one biotin, two or three conjugated biotin moieties per mAb, respectively.

(**C**) PALM measurements of 4% PFA-pretreated human T cells, which had not been decorated with any fluorescent probe, gave rise to no or only a few fluorescence signals detected within 10,000 frames for all three different color channels used for PS/PA fluorophore characterization.

(**D**) Classification of TCR CD3ζ- and LFA-1- distributions on 4% PFA-fixed T cells. Top: randomized distributions superimposed with experimentally determined blinking parameters of mEOS3.2 were simulated at densities of 38 (CD3ζ) and 45 (LFA-1) molecules $\mu m^{-2}$ (blue) for comparison with localization maps derived from PALM experiments involving CD3ζ-mEOS3.2 or anti-LFA-1-biotin decorated with mSAv*-cc-mEOS3.2* (magenta). Bottom: Ripley's K functions of CD3ζ-mEOS3.2 and anti-LFA-1-biotin-mSAv*-cc-mEOS3.2* (magenta) were compared to Ripley's K functions derived from 10 simulations of randomized distributions with corresponding molecular densities (blue, mean ± SD). Scale bars: 1 μm.

(**E**) Classification of TCRβ– and LFA-1- distributions on 4% PFA-fixed T cells. Top: randomized distributions superimposed with experimentally determined blinking parameters of CAGE635 were simulated at densities of 33 TCRβ and 25 LFA-1 molecules $\mu m^{-2}$ (blue) for comparison with localization maps derived from PALM experiments involving anti-TCRβ-biotin (mAb H57-biotin) visualized with mSAv*-c-CAGE635 and anti-LFA-1-biotin decorated with mSAv*-c-CAGE635 (magenta). Bottom: Ripley's K functions of TCRβ-biotin-mSAv*-c-CAGE635 and anti-LFA-1-biotin-mSAv*-c-CAGE635 (magenta) were compared to Ripley's K functions derived from 10 simulations of randomized distributions with corresponding molecular densities (blue, mean ± SD). Scale bars: 1 μm.

### Supplementary Figure 7

## A

```
1        10        20        30        40        50        60        70
```

MAEAGITGTW YNQLGSTFIV TAGADGALTG TYESAVGNAE SRYVLTGRYD SAPATDGSGT ALGWTVAWKN
NYRNAHSATT WSGQYVGGAE ARINTQWLLT SGTTE<u>A</u>NAWK STLVGHDTFT KVKPSAASGG LEVLFQGPEE
EEEE-

## B

```
1        10        20        30        40        50        60        70
```

**CATATG**GCTG AAGCTGGTAT CACCGGCACC TGGTACAACC AGCTGGGATC CACCTTCATC GTTACCGCTG
GTGCTGACGG TGCTCTGACC GGTACCTACG AATCCGCTGT TGGTAACGCT GAATCTAGAT ACGTTCTGAC
CGGTCGTTAC GACTCCGCTC CGGCTACCGA CGGTTCCGGA ACCGCTCTGG GTTGGACCGT TGCTTGGAAA
AACAACTACC GTAACGCTCA CTCCGCTACC ACCTGGTCTG GCCAGTACGT TGGTGGTGCT GAAGCTCGTA
TCAACACCCA GTGGTTGTTG ACCTCCGGCA CCACCGAA<u>GC C</u>AACGCGTGG AAATCCACCC TGGTTGGTCA
CGACACCTTC ACCAAAGTTA AACCGTCCGC TGCTTCCGGT GGTCTGGAAG TTCTGTTTCA AGGTCCGGAA
GAGGAAGAGG AAGAGTAATA A**AAGCTT**

**Supplementary Figure 7: Sequence of "alive" streptavidin subunit.**

(**A**) Protein sequence of the "alive" streptavidin subunit. We inserted a 3C protease cleavage site (blue) upstream of the poly glutamate tag (red). The alanine residue which has been substituted for a cysteine residue in the mSAv* construct is underlined.

(**B**) DNA sequence of the "alive" streptavidin subunit. The sequence for the 3C protease cleavage site is colored in blue, the sequence encoding the poly glutamate tag in red. The mutation resulting in A106C (GCC > TGC) is underlined. The construct was cloned into the pEt21a(+) expression vector with the 5´restriction enzyme NdeI and the 3´restriction enzyme HindIII (highlighted in bold).

### Supplementary Figure 8

## A

```
1        10          20          30          40          50          60          70
MAEAGITGTW YAQLGDTFIV TAGADGALTG TYEAAVGNAE SRYVLTGRYD SAPATDGSGT ALGWTVAWKN
NYRNAHSATT WSGQYVGGAE ARINTQWLLT SGTTEANAWK STLVGHDTFT KVKPSAASLE HHHHHH-
```

## B

```
1        10          20          30          40          50          60          70
CATATGGCTG AAGCTGGTAT CACCGGCACC TGGTACGCCC AGCTGGGAGA CACCTTCATC GTTACCGCTG
GTGCTGACGG TGCTCTGACC GGTACCTACG AAGCCGCTGT TGGTAACGCT GAATCTAGAT ACGTTCTGAC
CGGTCGTTAC GACTCCGCTC CGGCTACCGA CGGTTCCGGA ACCGCTCTGG GTTGGACCGT TGCTTGGAAA
AACAACTACC GTAACGCTCA CTCCGCTACC ACCTGGTCTG GCCAGTACGT TGGTGGTGCT GAAGCTCGTA
TCAACACCCA GTGGTTGTTG ACCTCCGGCA CCACCGAAGC TAACGCGTGG AAATCCACCC TGGTTGGTCA
CGACACCTTC ACCAAAGTTA AACCGTCCGC TGCTTCCCTC GAGCACCACC ACCACCACCA CTGA
```

**Supplementary Figure 8: Sequence of the "dead" streptavidin subunit.**

(**A**) Protein sequence of the "dead" streptavidin subunit. To arrive at mSAv*-3xHis$_6$, we extended the "dead" streptavidin subunit C-terminally with a 6 x histidine tag (marked in green).

(**B**) DNA sequence encoding the "dead" streptavidin. The sequence encoding the 6 x histidine tag is marked in green. The construct was cloned into the pEt21a(+) expression vector with the 5´restriction enzyme NdeI and the 3´restriction enzyme XhoI (highlighted in bold).

### Supplementary Figure 9

## A

```
1         10        20        30        40        50        60        70
MSKGAELFTG IVPILIELNG DVNGHKFSVS GEGEGDATYG KLTLKFICTT GKLPVPWPTL VATLSYGVQC
FSRYPDHMKQ HDFFKSAMPE GYIQERTIFF EDDGNYKTRA EVKFEGDTLV SRIELTGTDF KEDGNILGNK
MEYNYNATNV YIVADKARNG IKVNFKVRHN IKDGSVQLAD HYQQNTPIGD GPVLLPDNHY LSTQSALSKD
PNEKRDHMIY LEFVTAAAIT HGMDELYKGG GSGLNDIFEA QKIEWHEGSG GLEVLFQGPH HHHHHHHHHH
H-
```

## B

```
1         10        20        30        40        50        60        70
CATATGAGCA AGGGCGCCGA GCTGTTCACC GGCATCGTGC CCATCCTGAT CGAGCTGAAT GGCGATGTGA
ATGGCCACAA GTTCAGCGTG AGCGGCGAGG GCGAGGGCGA TGCCACCTAC GGCAAGCTGA CCCTGAAGTT
CATCTGCACC ACCGGCAAGC TGCCTGTGCC CTGGCCCACC CTGGTGGCCA CCCTGAGCTA CGGCGTGCAG
TGCTTCTCAC GCTACCCCGA TCACATGAAG CAGCACGACT TCTTCAAGAG CGCCATGCCT GAGGGCTACA
TCCAGGAGCG CACCATCTTC TTCGAGGATG ACGGCAACTA CAAGACGCGC GCCGAGGTGA AGTTCGAGGG
CGATACCCTG GTGAGTCGCA TCGAGCTCAC CGGCACTGAT TTCAAGGAGG ATGGCAACAT CCTGGGCAAT
AAGATGGAGT ACAACTACAA CGCCACCAAT GTGTACATCG TGGCCGACAA GGCCAGGAAT GGCATCAAGG
TGAACTTCAA GGTCCGCCAC AACATCAAGG ATGGCAGCGT GCAGCTGGCC GACCACTACC AGCAGAATAC
CCCCATCGGC GATGGCCCTG TGCTGCTGCC CGATAACCAC TACCTGTCCA CCCAGAGCGC CCTGTCCAAG
GACCCCAACG AGAAGCGCGA TCACATGATC TACCTCGAGT TCGTGACCGC CGCCGCCATC ACCCACGGCA
TGGATGAGCT GTACAAGGGT GGCGGATCCG GCCTGAACGA TATTTTTGAA GCGCAGAAAA TTGAATGGCA
TGAAGGTAGC GGCGGTCTGG AAGTTCTGTT TCAAGGTCCG CATCACCATC ACCATCACCA CCACCATCAT
CATCACTAAT AAAAGCTT
```

**Supplementary Figure 9: Sequence of PS-CFP2-biotin.**

**(A)** Protein sequence of the PS-CFP2-AVI-3C-$His_{12}$ construct with the PS-CFP2 protein chain marked in black, the BirA ligase recognition sequence in violet, the 3C protease cleavage site in blue, the poly-histidine tag in red and the linker sequences in green. The underline serine residue had been substituted for a cysteine residue.

**(B)** DNA sequence of the PS-CFP2-AVI-3C-$His_{12}$ construct. The PS-CFP2 gene was extended with linker sequences (green), a BirA ligase recognition site (violet), a 3C protease cleavage site (blue) and a poly-histidine tag (red). The nucleotide substitution (AGC > TGC) encoding for a cysteine residue is underlined. The construct was cloned into the pEt21a(+)expression vector with the 5´restriction enzyme NdeI and the 3´restriction enzyme HindIII (in bold font).

### Supplementary Figure 10

## A

```
1         10                20                30                40                50                60                70
MSAIKPDMKI KLRMEGNVNG HHFVIDGDGT GKPFEGKQSM DLEVKEGGPL PFAFDILTTA FHYGNRVFAK
YPDNIQDYFK QSFPKGYSWE RSLTFEDGGI CNARNDITME GDTFYNKVRF YGTNFPANGP VMQKKTLKWE
PSTEKMYVRD GVLTGDIEMA LLLEGNAHYR CDFRTTYKAK EKGVKLPGAH FVDHCIEILS HDKDYNKVKL
YEHAVAHSGL PDNARRGSGL NDIFEAQKIE WHEGSGGLEV LFQGPHHHHH HHHHHHH--
```

## B

```
1         10                20                30                40                50                60                70
CATATGAGTG CGATTAAGCC AGACATGAAG ATCAAACTCC GTATGGAAGG CAACGTAAAC GGGCACCACT
TTGTGATCGA CGGAGATGGT ACAGGCAAGC CTTTTGAGGG AAAACAGAGT ATGGATCTTG AAGTCAAAGA
GGGCGGACCT CTGCCTTTTG CCTTTGATAT CCTGACCACT GCATTCCATT ACGGCAACAG GGTATTCGCC
AAATATCCAG ACAACATACA AGACTATTTT AAGCAGTCGT TTCCTAAGGG GTATTCGTGG GAACGAAGCT
TGACTTTCGA AGACGGGGGC ATTTGCAACG CCAGAAACGA CATAACAATG GAAGGGGACA CTTTCTATAA
TAAAGTTCGA TTTTATGGTA CCAACTTTCC CGCCAATGGT CCAGTTATGC AGAAGAAGAC GCTGAAATGG
GAGCCCTCCA CTGAGAAAAT GTATGTGCGT GATGGAGTGC TGACGGGTGA TATTGAGATG GCTTTGTTGC
TTGAAGGAAA TGCCCATTAC CGATGTGACT TCAGAACTAC TTACAAAGCT AAGGAGAAGG GTGTCAAGTT
ACCAGGCGCC CACTTTGTGG ACCACTGCAT TGAGATTTTA AGCCATGACA AAGATTACAA CAAGGTTAAG
CTGTATGAGC ATGCTGTTGC TCATTCTGGA TTGCCTGACA ATGCCAGACG AGGATCCGGC CTGAACGATA
TTTTTGAAGC GCAGAAAATT GAATGGCATG AAGGTAGCGG CGGTCTGGAA GTTCTGTTTC AAGGTCCGCA
TCACCATCAC CATCACCACC ACCATCATCA TCACTAATAA AAGCTT
```

**Supplementary Figure 10: Sequence of mEOS3.2-biotin.**

**(A)** Protein sequence of the mEOS3.2-AVI-3C-His$_{12}$ construct. The mEOS3.2 protein chain is marked in black, the BirA ligase recognition sequence in violet, the 3C protease cleavage site in blue, the poly-histidine tag in red and the linker sequences in green.

**(B)** DNA sequence of the mEOS3.2-AVI-3C-His$_{12}$ construct. The mEOs3.2 gene was extended with linker sequences (green), a BirA ligase recognition site (violet), a 3C protease cleavage site (blue) and a poly-histidine tag (red). The construct was cloned into the pEt21a(+)expression vector with the 5´restriction enzyme NdeI and the 3´restriction enzyme BamHI (in bold font).

**Supplementary Table 1:**

Two-sample Kolmogorov-Smirnov test to determine the p-value for the hypothesis that two different data sets are from the same continuous distribution. Distributions of all indicated imaging conditions were tested against each other. This was done for all investigated fluorophores and for all four parameters: (i) the total number of detections (***N***), (ii) the number of off-gaps (***g***), (iii) the duration of each emission burst ($\boldsymbol{t_{on}}$), and (iv) the duration of each off-gap ($\boldsymbol{t_{off}}$). Significant p-values ($p \leq 0.05$) are indicated in bold font. Laser power density is given in kW $cm^{-2}$ and abbreviated with kW in all tables.

### Supplementary Table 1

PSCFP-2

| | | 2 ms / 3.0 kW | 15 ms / 0.4 kW | 2 ms / 3.0 kW + GSH |
|---|---|---|---|---|
| number of detections | 2 ms / 3.0 kW + 4% PFA | 0.107 | 0.680 | 0.174 |
| | 2 ms / 3.0 kW | - | 0.134 | 0.268 |
| | 15 ms / 0.4 kW | - | - | 0.113 |
| number of gaps | 2 ms / 3.0 kW + 4% PFA | 0.562 | 0.991 | 0.564 |
| | 2 ms / 3.0 kW | - | 0.484 | 0.522 |
| | 15 ms / 0.4 kW | - | - | 0.326 |
| $t_{on}$ | 2 ms / 3.0 kW + 4% PFA | 0.371 | 0.955 | 0.475 |
| | 2 ms / 3.0 kW | - | 0.243 | 0.912 |
| | 15 ms / 0.4 kW | - | - | 0.238 |
| $t_{off}$ | 2 ms / 3.0 kW + 4% PFA | **1.32E-24** | **6.94E-08** | **0.005** |
| | 2 ms / 3.0 kW | - | **1.58E-07** | 0.186 |
| | 15 ms / 0.4 kW | - | - | **1.97E-04** |

mEOS3.2

| | | 2 ms / 4.0 kW | 15 ms / 0.57 kW |
|---|---|---|---|
| number of detections | 2 ms / 4.0 kW + 4% PFA | 0.456 | 0.451 |
| | 2ms / 4.0 kW | - | 0.993 |
| number of gaps | 2 ms / 4.0 kW + 4% PFA | 0.997 | 0.611 |
| | 2ms / 4.0 kW | - | 0.655 |
| $t_{on}$ | 2 ms / 4.0 kW + 4% PFA | 0.998 | 0.106 |
| | 2ms / 4.0 kW | - | 0.102 |
| $t_{off}$ | 2 ms / 4.0 kW + 4% PFA | 0.393 | 0.460 |
| | 2ms / 4.0 kW | - | 0.425 |

PA Janelia Fluor 549

| | | 2 ms / 4.0 kW | 15 ms / 0.53 kW |
|---|---|---|---|
| number of detections | 2 ms / 4.0 kW + 4% PFA | 0.562 | **6.04E-06** |
| | 2ms / 4.0 kW | - | **0.006** |
| number of gaps | 2 ms / 4.0 kW + 4% PFA | 0.483 | **1.15E-05** |
| | 2ms / 4.0 kW | - | **0.032** |
| $t_{on}$ | 2 ms / 4.0 kW + 4% PFA | 1.000 | 0.985 |
| | 2ms / 4.0 kW | - | 1.000 |
| $t_{off}$ | 2 ms / 4.0 kW + 4% PFA | 0.997 | **0.006** |
| | 2ms / 4.0 kW | - | **0.030** |

Abberior CAGE 635

| | | 2 ms / 4.0 kW | 15 ms / 0.53 kW |
|---|---|---|---|
| number of detections | 2 ms / 4.0 kW + 4% PFA | 0.162 | 0.053 |
| | 2ms / 4.0 kW | - | **0.001** |
| number of gaps | 2 ms / 4.0 kW + 4% PFA | **0.047** | **0.002** |
| | 2ms / 4.0 kW | - | **1.67E-04** |
| $t_{on}$ | 2 ms / 4.0 kW + 4% PFA | 0.959 | 0.421 |
| | 2ms / 4.0 kW | - | 0.403 |
| $t_{off}$ | 2 ms / 4.0 kW + 4% PFA | 0.899 | 0.708 |
| | 2ms / 4.0 kW | - | 0.979 |

**Supplementary Table 2:**

Two-sample Kolmogorov-Smirnov test to determine the p-value for the hypothesis that two different data sets are from the same continuous distribution. Distributions of all indicated PA/PS-fluorophores were tested against each other, for all imaging conditions and for all four parameters: (i) the total number of detections (***N***), (ii) the number of off-gaps (***g***), (iii) the duration of each emission burst ($t_{on}$), and (iv) the duration of each off-gap ($t_{off}$). Significant p-values ($p \leq 0.05$) are indicated in bold font. Laser power density is given in kW $cm^{-2}$ and abbreviated with kW in all tables.

### Supplementary Table 2

2 ms illumination and 4 % PFA

| | | mEOS3.2 | PA Janelia Fluor 549 | Abberior CAGE 635 |
|---|---|---|---|---|
| number of detections | PSCFP-2 | **0.009** | **4.68E-09** | **3.11E-35** |
| | mEOS3.2 | - | 0.699 | **9.50E-11** |
| | PA Janelia Fluor 549 | - | - | **1.39E-22** |
| number of gaps | PSCFP-2 | 0.650 | **5.59E-04** | **9.76E-12** |
| | mEOS3.2 | - | 0.693 | **1.70E-05** |
| | PA Janelia Fluor 549 | - | - | **2.44E-12** |
| $t_{on}$ | PSCFP-2 | **0.046** | 1.000 | **1.62E-70** |
| | mEOS3.2 | - | **0.046** | **1.62E-17** |
| | PA Janelia Fluor 549 | - | - | **1.61E-46** |
| $t_{off}$ | PSCFP-2 | **1.93E-19** | **9.69E-15** | **3.51E-66** |
| | mEOS3.2 | - | **4.74E-04** | 0.106 |
| | PA Janelia Fluor 549 | - | - | **4.61E-08** |

2 ms illumination

| | | mEOS3.2 | PA Janelia Fluor 549 | Abberior CAGE 635 |
|---|---|---|---|---|
| number of detections | PSCFP-2 | **0.023** | **0.004** | **1.01E-45** |
| | mEOS3.2 | - | 0.299 | **3.03E-11** |
| | PA Janelia Fluor 549 | - | - | **1.34E-15** |
| number of gaps | PSCFP-2 | 0.604 | 0.235 | **6.41E-26** |
| | mEOS3.2 | - | 0.998 | **6.40E-07** |
| | PA Janelia Fluor 549 | - | - | **3.00E-10** |
| $t_{on}$ | PSCFP-2 | **0.005** | 0.160 | **2.17E-88** |
| | mEOS3.2 | - | 0.084 | **1.45E-13** |
| | PA Janelia Fluor 549 | - | - | **7.24E-34** |
| $t_{off}$ | PSCFP-2 | **1.17E-24** | **4.65E-19** | **9.53E-112** |
| | mEOS3.2 | - | 0.107 | 0.181 |
| | PA Janelia Fluor 549 | - | - | **1.23E-06** |

15 ms illumination

| | | mEOS3.2 | PA Janelia Fluor 549 | Abberior CAGE 635 |
|---|---|---|---|---|
| number of detections | PSCFP-2 | **0.023** | **0.004** | **1.01E-45** |
| | mEOS3.2 | - | 0.299 | **3.03E-11** |
| | PA Janelia Fluor 549 | - | - | **1.34E-15** |
| number of gaps | PSCFP-2 | 0.604 | 0.235 | **6.41E-26** |
| | mEOS3.2 | - | 0.998 | **6.40E-07** |
| | PA Janelia Fluor 549 | - | - | **3.00E-10** |
| $t_{on}$ | PSCFP-2 | **0.005** | 0.160 | **2.17E-88** |
| | mEOS3.2 | - | 0.084 | **1.45E-13** |
| | PA Janelia Fluor 549 | - | - | **7.24E-34** |
| $t_{off}$ | PSCFP-2 | **1.17E-24** | **4.65E-19** | **9.53E-112** |
| | mEOS3.2 | - | 0.107 | 0.181 |
| | PA Janelia Fluor 549 | - | - | **1.23E-06** |
